# The Optimal Retinal Locus for High-Resolution Vision in Space and Time

**DOI:** 10.1101/2025.04.30.650879

**Authors:** Josselin Gautier, Norick R. Bowers, Martin S. Banks, Austin Roorda

## Abstract

Humans exhibit machine-like eye-movement behavior in space and time while performing challenging visual resolution tasks. Fewer microsaccades occur as stimulus presentation is imminent. Drifts and microsaccades combine to confine the landing location of an anticipated visual stimulus to a tiny retinal region: the preferred retinal locus (PRL). We find that this location confers the best visual acuity despite it being offset from the anatomical fovea (the location of maximum cone density). We also find that acuity is best when the last microsaccade occurs 400msec or longer before stimulus presentation. The machine-like eye movements are involuntary and not perceived. Our findings thus reveal a highly evolved oculomotor system such that gaze direction during fixation is rarely far enough from the PRL to cause a decline in visual resolution.

**Teaser:** Humans exhibit machine-like eye-movement behavior during fine visual tasks that rarely deviates from an optimal retinal location.

## Introduction

Most animals have fairly constant receptor density across the retina (*5–7*). But many, including humans, are different in that they have a very specialized area in the retina—the fovea—with much more tightly packed photoreceptors than other parts of the retina (*8–10*). The human fovea has no rods and few short-wavelength (S) cones (*10,11*). The fovealar long- and medium-wavelength (L & M) cones each make connections to at least two ganglion cells (*12, 13*) and have a magnified representation in the visual cortex (*14, 15*). These specializations enable the resolution of much finer detail with the fovea than with other parts of the retina (*3, 16–20*). In Fig. 1A, we plot data from four studies that measured acuity at or very close to the line of sight. Acuity falls off precipitously with increasing eccentricity. The fall off is similar for letter and grating acuity (*2, 3*), but is steepest for hyperacuity tasks (*21, 22*). Thus, it is very well established that the ability to resolve fine detail is much better for stimuli falling on the fovea than for stimuli falling on other parts of the retina.

**Figure 1:**
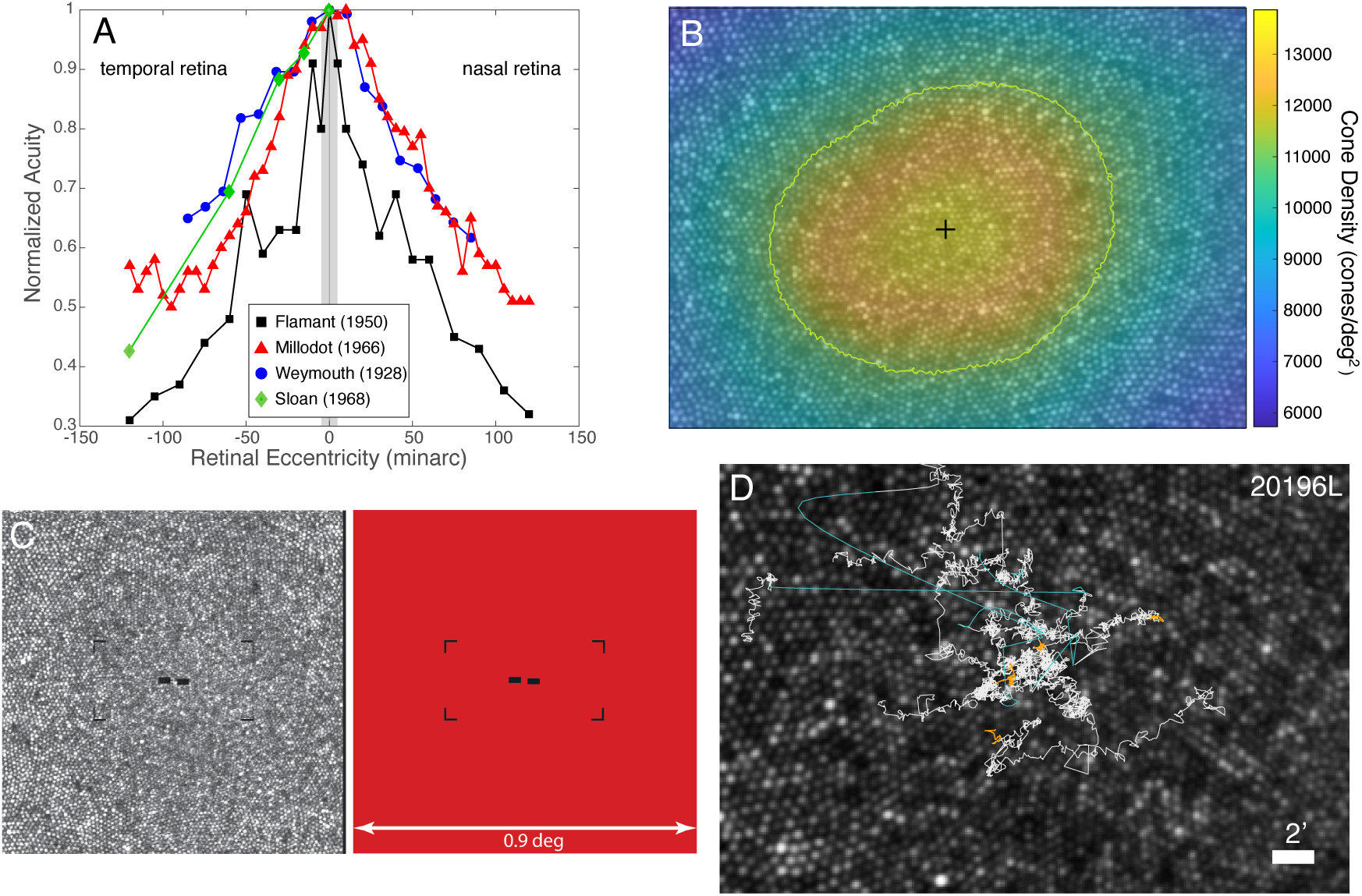
Introduction and methods. **A)** Measurements of visual acuity as a function of eccentricity are plotted for four previous studies that made measurements near the line of sight. The data have been normalized, so the highest acuities have a value of 1 on the ordinate. Data from the temporal and nasal retina are on the left and right, respectively. Black squares represent data from Flamant (*1*): Two-dot separation acuities from one subject. Red triangles represent data from Millodot (*2*): Landolt-C acuities averaged from three subjects. Blue circles are data from Weymouth and colleagues (*3*): grating acuities from one subject. Green diamonds are from Sloan (*4*): Landolt-C acuities at 1.5 log millilamberts from one subject. The gray rectangle represents eccentricities spanning +/-10 minarc from the line of sight, which is the region we focused on in the current study. **B)** Cone mosaic for one subject. Density (cones per square deg) is represented by color as indicated by the color bar. The green contour represents where density falls to 80% of the peak. The black cross represents the cone density centroid (CDC), which is the density-weighted centroid within the 80% contour. **C)** The left panel shows one raw AOSLO frame. The fixation guides and Vernier stimulus are generated by turning off the same 680nm laser that is used to record the image. Thus the stimulus and fixation guides are encoded directly into the AOSLO frame. The right panel shows how the raster appears to the subject. Field size is 0.9×0.9^◦^, fixation guides are 25×12.5minarc apart, and each bar of the stimulus is 0.95×1.9minarc, separated by 0.95minarc. **D)** Traces of target position on the retina for five consecutive trials in a representative subject. The stimulus was presented for 34msec. Its position during that time is indicated by the orange traces. We also plot where it would have fallen on the retina across the whole 2-sec trial. White traces represent drifts. Cyan traces represent microsaccades. Scale bar indicates 2minarc.

Advances in retinal-imaging instrumentation have enabled researchers to precisely determine the part of the retina people use when asked to fixate a small target. Interestingly, people consistently place the target near the anatomical fovea (defined here as the retinal location with highest cone density), but on average about 4–10minarc away (*23–25*). This location is called the preferred retinal locus or PRL. The small PRL displacement is consistent over long periods of time (*25, 26*) and across different visual tasks (*27, 28*). Its position relative to the anatomical fovea differs across individuals, but is frequently displaced upward in retinal coordinates from the center of the fovea (*23, 25, 28*). It is very important to note that previous studies of visual resolution across the retina have assumed that observers fixate with the foveal center when given a fixation target (*3, 16, 18*). Thus, they assume that the retinal eccentricity of the stimulus is its angular distance from the center of the fovea. But the recent findings that fixation is consistently displaced from the center mean that this assumption is false.

When an observer tries to maintain steady gaze on a small target, the eyes still move. These fixational eye movements are characterized by slow movements, or drifts, interspersed with fast movements that are small saccades, or microsaccades (*27*). In previous measurements of acuity as a function of retinal eccentricity, drifts and microsaccades would have caused the stimulus to fall at different positions on the retina despite the observer’s attempt to maintain steady gaze. In other words, uncontrolled fixational movements would cause uncertainty about the actual position of the stimulus on the retina. This uncertainty in turn should smooth out a plot of acuity vs eccentricity, leading to the misleading impression that acuity varies less around the center of gaze than it actually does (*29*). This dilemma is potentially solved by stabilizing the acuity target on the retina (*2, 29*) but, without imaging the retina, one still cannot know where the stabilized stimulus fell relative to the anatomical fovea; if it falls on different positions from trial to trial, one still expects to observe a smoothed plot of acuity vs eccentricity, again leading to the possibly false impression that acuity plateaus near the fovea.

Researchers have recently used high-resolution retinal imaging combined with visual psychophysics to better investigate how visual performance varies with retinal eccentricity. Specifically, they investigated how performance varies for stimuli on the PRL or systematically displaced from it. Domdei and colleagues (*30*) presented small spots at different prespecified locations on the retina. They used stabilized imagery to position stimuli precisely at the PRL or 6 or 12minarc from it. They reported that detectability was essentially constant across the region tested: specifically, that detectability was no better at the PRL than at the locus of peak cone density (*30*). Ratnam (*31*) conducted a letter-acuity task for stabilized stimuli presented at the PRL and up to 10minarc from it and found no drop-off in that region of the retina. Witten and colleagues (*32*) argued that fixational movements serve to direct the fixated image toward retinal locations with higher cone densities, but they did not show that these higher density regions in the same individuals conferred greater acuity. In summary, using retinal imaging in tandem with either accurate recording or direct control of the stimulated region, researchers have been unable to determine whether or not visual sensitivity and acuity plateau across the small retinal region surrounding the PRL and peak cone density.

It has been argued that the incessant movement of the eyes during fixation should impair visual resolution, much like a shaky camera produces a blurred photograph (*33, 34*). But it has also been argued that the movements are beneficial to high-resolution vision for two reasons; first, because the eyes’ motion spreads stimulus features over many photoreceptors thereby effectively increasing the number of samples within a temporal integration window (*35–37*) and second, because the motion reshapes the spatiotemporal properties by equalizing or “whitening” spatial energy across the temporal domain (*38*). Related to this, we know that fixational eye movements differ when observers perform different tasks. For example, drifts become slower and more curved when performing an acuity task compared to simply maintaining fixation on a small target (*29*), and microsaccades are less frequent in an acuity task (*28*). From such data, it has been concluded that fixational eye movements are fine-tuned during resolution tasks so that the resulting positioning of the upcoming stimulus on the PRL and the reduced motion of the retinal image help maximize acuity (*28, 29*).

To investigate these issues further, we examined natural fixations during a cadenced visualacuity task. We wanted to know how performance is affected by the position of the stimulus on the retina and by the variation in drifts and microsaccades before, during, and after the presentation of a brief stimulus. We emphasize that the eye movements were voluntary and the retinal images unstabilized so that the position and motion of the stimulus on the retina was natural.

## Results

Essential parts of the methods are described next. Full details are in the Materials and Methods section. Data were collected for five subjects. Each subject’s data consisted of 100 44-sec videos, each containing 21 trials. Fixation guides (Fig. 1C) were presented on the left and right, upper, and lower corners of the stimulus field. Subjects were asked to maintain fixation in the middle of the guides. On the 60th frame of each trial, the Vernier stimulus (Fig. 1C) was presented at the center of the guides for two frames. The subject then indicated whether the left bar was higher or lower than the right bar, yielding 2100 responses for each subject. The complete eye-motion trace was computed for each video and was used to determine the trajectory that the center of the fixation guides traced on the retina. In this way the exact retinal location of the Vernier stimulus could be determined for each trial. Fig. 1D shows eye traces for five consecutive trials overlaid onto a high-resolution master retinal image for one subject. The traces are color-coded to show drifts (white) and saccades (cyan), and the trace when the stimulus was being presented (orange). All the data from the complete eye-motion traces and 2100 trials were registered to a master retinal image. This comprised the master data set from which all analyses were drawn.

The motion trajectories were annotated to identify epochs of drift, the start and end points of microsaccades, and the time and location of the stimulus. This enabled a computation of the distribution of the retinal locations of the center of the fixation guides at all time points during the cadenced task. Fig. 2A plots these distributions on the master retinal image as isodensity contours that encompass 68% of the data points for the entire trajectory (gray), the saccade start points (blue), saccade end points (magenta), and the retinal locations where the stimulus landed (orange). We call the stimulus landing locations the functional PRL or fPRL. The peak of the kernel density distribution for each respective scatter plot of data points is indicated by a symbol of the same color. The areas of the isodensity contours (ISOA) encompassing 68% of the scatter plots of the data points are provided in Table 1.

**Figure 2:**
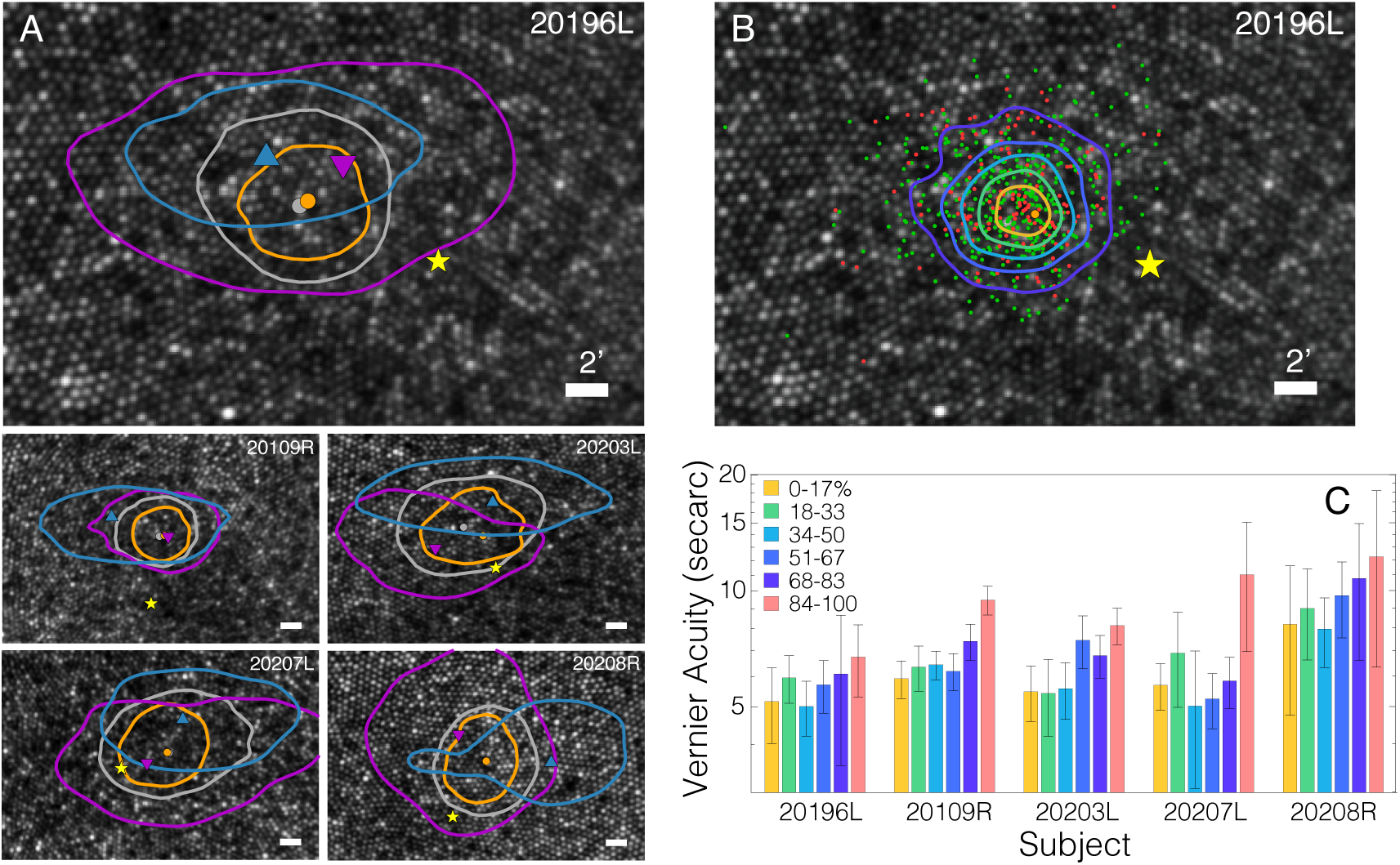
PRLs, ISO contours, and performance. **A)** PRLs and ISO contours from all 5 subjects. Gray dots and contours are the PRL and ISOA contours from the entire distribution of target locations for the 2sec trials. Orange dots and contours are the PRL and corresponding ISOA computed from the 34msec when the target was presented. Purple triangles and contours are computed from the starting position of each microsaccade: the Saccade Starting Position. Blue triangles and contours are computed from the landing position of each microsaccade: the Saccade Landing Position. These two positions are where the stimulus would have fallen if it were presented when the saccade started or ended. Yellow stars are the location of the CDC. Scale bars indicate 2minarc. **B)** Stimulus positions and performance. The position of the stimulus across trials is plotted for a representative observer. Green dots represent the retinal loci of the stimulus when the observer responded correctly. Red dots are for incorrect responses. The yellow star indicates the CDC. Orange, green, cyan, blue, and purple curves are the ISO contours that contain 17, 33, 50, 67, and 83% of the responses, respectively. Scale bar represents 2minarc. **C)** Acuity thresholds based on where the stimulus fell. Different sets of histograms are for different observers. The colors are, respectively, thresholds based on the 17% of positions falling closest to the PRL (yellow), 18–33% (green), 34–50% (cyan), 51-67% (blue), 68–83% (purple), and 84–100% (pink). Error bars are standard errors.

**Table 1:** Subject data. All averages are arithmetic averages except for Vernier acuity which was converted to logarithms before averaging. Nyq lim (c/d) refers to Nyquist sampling limit of the cone mosaic in cycles per degree.

| Subject ID | Tested Eye | Vernier Acuity (secarc) | ISOA (minarc<sup>2</sup>) |  |  |  | fPRL-CDC x,y(minarc) | Nyq lim (c/d) |  |
| --- | --- | --- | --- | --- | --- | --- | --- | --- | --- |
|  |  |  | TotPRL | fPRL | SacStart | SacEnd |  | fPRL | CDC |
| 20196 | L | 6.29 ± 0.27 | 55.18 | 25.68 | 176.97 | 69.95 | 5.87 | 67.0 | 68.2 |
| 20109 | R | 7.46 ± 0.31 | 34.62 | 22.99 | 69.65 | 101.96 | 6.23 | 65.3 | 68.8 |
| 20203 | L | 6.98 ± 0.32 | 102.29 | 51.70 | 138.13 | 132.49 | 2.94 | 64.0 | 64.0 |
| 20207 | L | 7.40 ± 0.36 | 98.64 | 50.67 | 229.59 | 141.47 | 3.87 | 69.7 | 70.3 |
| 20208 | R | 10.76 ± 0.55 | 77.00 | 44.04 | 231.21 | 120.76 | 6.59 | 60.4 | 62.8 |
| AVG |  | 7.64 | 73.54 | 39.02 | 169.11 | 113.33 | 5.10 | 65.3 | 66.8 |

A key observation is that the fPRL is the smallest of the four, suggesting that the entire trajectory of the eye motion during the task serves to confine the landing position of the anticipated stimulus to a small retinal area. The ISOA for the fPRL is 39.02minarc^2^ on average, which is about two times smaller than the overall ISOA. We note that the overall ISOA is the more conventional definition for the PRL (*39*). The ISOAs for the saccade starting and ending points are larger, and their contour plots are elongated horizontally, and generally displaced vertically from the overall and functional ISOA contours. These tendencies are consistent with earlier reports of larger and more frequent horizontal microsaccades (*40, 41*) along with a slight tendency for microsaccades to have an upward vertical component (*28*).

We define the location of the anatomical fovea as the location of the cone density centroid (CDC), which corresponds to the centroid of the region of peak density (*25*) (Fig. 1B). The CDC is also labeled on each master retinal image. The fPRL is displaced from the CDC by 5.10minarc on average and is above the CDC in all cases. These two observations are consistent with previous reports (*24,25*). The ISOA contour containing 68% of the stimulus landing points does not include the CDC in any of our subjects.

### Spatial Analysis

The Vernier-acuity thresholds were derived from the subjects’ responses in a two-alternative, forced-choice procedure with the method of constant stimuli. We fit the psychometric data with logistic functions using a maximum-likelihood criterion (*42*) and computed the acuity estimates from the fitted curves. The acuities were 7.64secarc on average, which is consistent with high-quality measures of Vernier acuity (*21,22,43*). The fact that the acuities were very high confirms the effectiveness of the stimulus-delivery system, the fixation guides, and the geometry of the stimuli. The individual and average acuity values are provided in Table 1.

We next computed Vernier acuity as a function of where the stimulus fell on the retina trial by trial. Fig. 2 shows the results. Fig. 2B displays, for a representative subject, the retinal location of the stimulus on trials in which the subject’s response was correct (green dots) and trials in which the response was incorrect (red dots). It also shows contours that contain the 17% of positions that are closest to the functional PRL (orange), 18–33% of the closest (green), 34– 50% (cyan), 51–67% (blue), and 68–83% (purple). There were 357 observations per subject in each region, including 84–100% (no contour). Fig. S1 shows the data for the other four subjects. Fig. 2C plots for each subject the acuity thresholds based on the data in each of the six regions. Lower thresholds (*i.e*, higher acuities) were obtained when the stimulus was closer to the PRL. The decrease in acuity with increasing distance from the PRL was highly reliable statistically (1-way ANOVA; F = 42.07, p*<*0.001). Paired-comparison tests show that the effect is due to the fall-off in performance for points 84% or farther from the PRL (post-hoc t-tests, p*<*0.001, Holm correction). All other pairwise comparisons did not reach statistical significance. Thus, during this cadenced acuity task, fixational eye movements placed the stimulus in a non-optimal position only about 17% of the time, leading to a moderate acuity loss of 34%.

We next asked if acuity for stimuli near the PRL is measurably better than acuity at or near the CDC. We did this by computing percent-correct response when the stimulus fell within 1.5minarc of the PRL relative to stimuli falling within that distance from the CDC. There were, of course, many more observations near the PRL because that location is by definition where stimuli were most likely to fall. Percent correct within 1.5minarc of the PRL (averaged across subjects) was 0.831 (STD = 0.041). Percent correct for the same distance from the CDC was 0.853 (0.027). These values did not differ statistically (t test: p = 0.398). We also asked whether acuity was better for points within 1.5minarc of the saccade landing position and found that it was not. Thus, Vernier acuity is essentially constant within a small region in the center of the retina even when subjects make natural fixational eye movements that position and move the stimulus on the retina.

### Temporal analysis

Subjects suppressed microsaccades prior to, during, and following the stimulus. Fig. 3A plots, for a representative subject, the frequency of microsaccades over the 2-sec duration of a trial. The time of stimulus presentation is represented by the gray bar and the saccade rate during correct and incorrect trials by the green and red lines, respectively. The microsaccade rate was slightly lower during stimulus presentation on correct trials than on incorrect ones. All subjects exhibited similar behavior (see Fig. S2): They reduced the number of microsaccades as the time of stimulus presentation approached and continued to do so for about 400msec after extinction of the stimulus. This was followed by a return to a higher saccade rate as they provided their response and awaited the next stimulus presentation. Surprisingly, the greatest suppression occurred about 200–400msec after the stimulus was extinguished.

**Figure 3:**
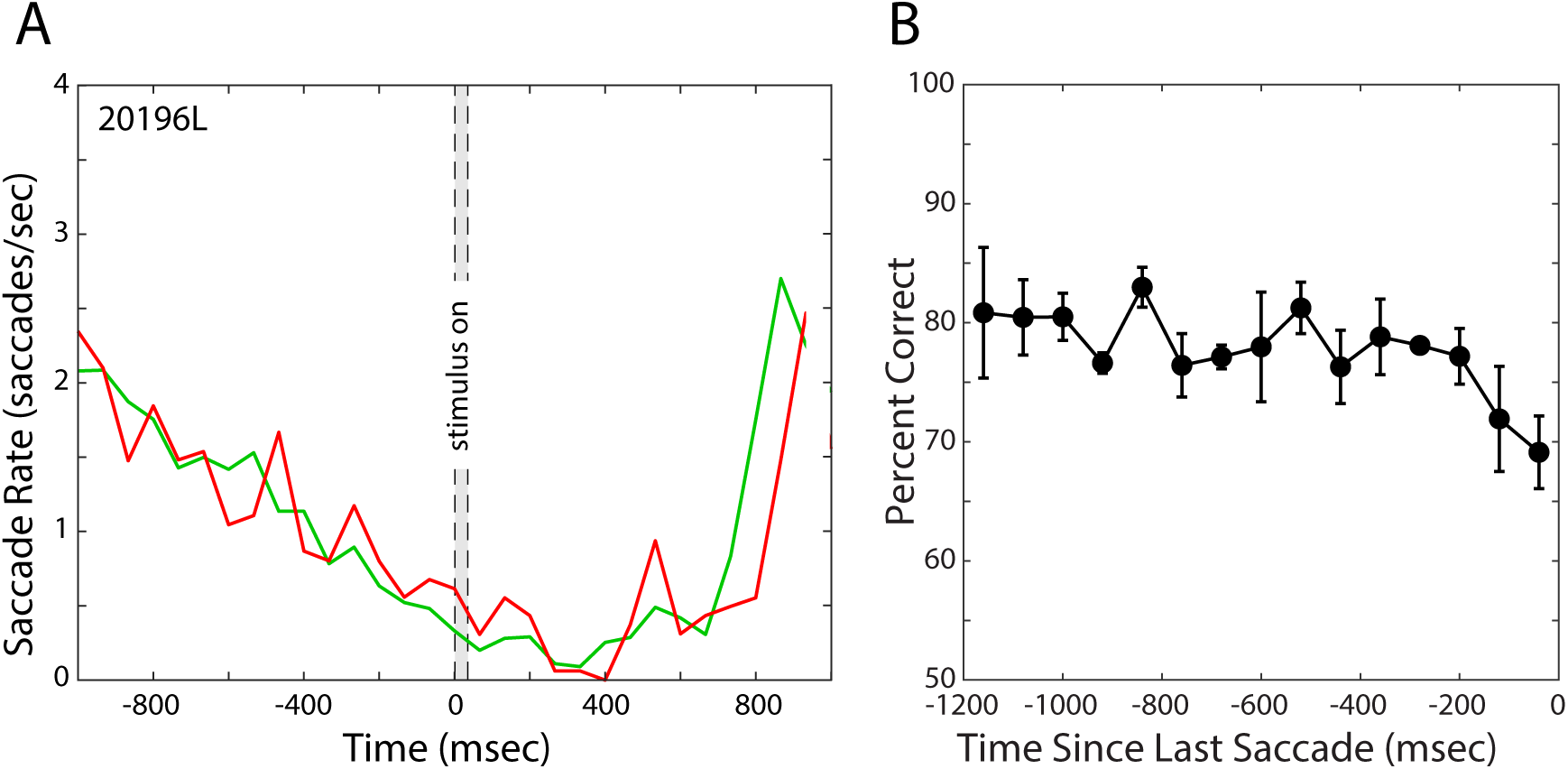
Saccade rate and performance. **A)**. Saccade rate for a representative subject over time. The average number of saccades per second is plotted as a function of time. Stimulus onset is at time 0. The gray bar represents the time the stimulus was presented. The green and red lines represent saccade rate during trials in which the observer responded correctly and incorrectly. **B)**. Last saccades and performance. Percent-correct performance in the acuity task is plotted as a function of how much time the last microsaccade preceded stimulus presentation. The data have been averaged across the 5 subjects. Each data point is the percent correct for 80msec bins centered on the time values on the abscissa. Percent correct was calculated excluding trials with Vernier offsets of +/-18secarc because performance was at ceiling for those trials. Error bars are standard errors of the mean.

We also examined when the last microsaccade occurred before the stimulus was presented and how that timing affected performance. The last microsaccades usually occurred 300–600msec before the stimulus appeared. Fig. 3B plots percent correct as a function of the time of the last microsaccade. It shows how that timing affected performance in the acuity task. There was a fairly consistent change in percent correct as a function of lag with shorter lags being associated with lower percent correct. For lags of 520msec and less, there was a marginally significant trend for poorer performance with shorter lags (ANOVA, F = 2.098, p = 0.085). Thus making a saccade roughly 400msec or less before stimulus onset hinders performance in a resolution task.

### Spatiotemporal Analysis

The consistency of fixation strategy during each cadenced trial is illustrated by Fig. 4. Fig. 4A plots the ISOA over the 2-sec duration of all trials for a representative subject. The data have been averaged across all 2100 trials. The green and red contours represent the ISOAs for trials in which the subject responded correctly and incorrectly, respectively. The ISOA becomes systematically smaller during the 1sec before stimulus onset. It is slightly smaller for correct than for incorrect trials. The area remains small for up to 400msec after the stimulus is extinguished, and then becomes much larger as the subject provided their response and prepared for the next trial. Every subject exhibited this general pattern (see Fig. S3). The change in ISOA is largely consistent with the suppression of microsaccades (Fig. 3A).

**Figure 4:**
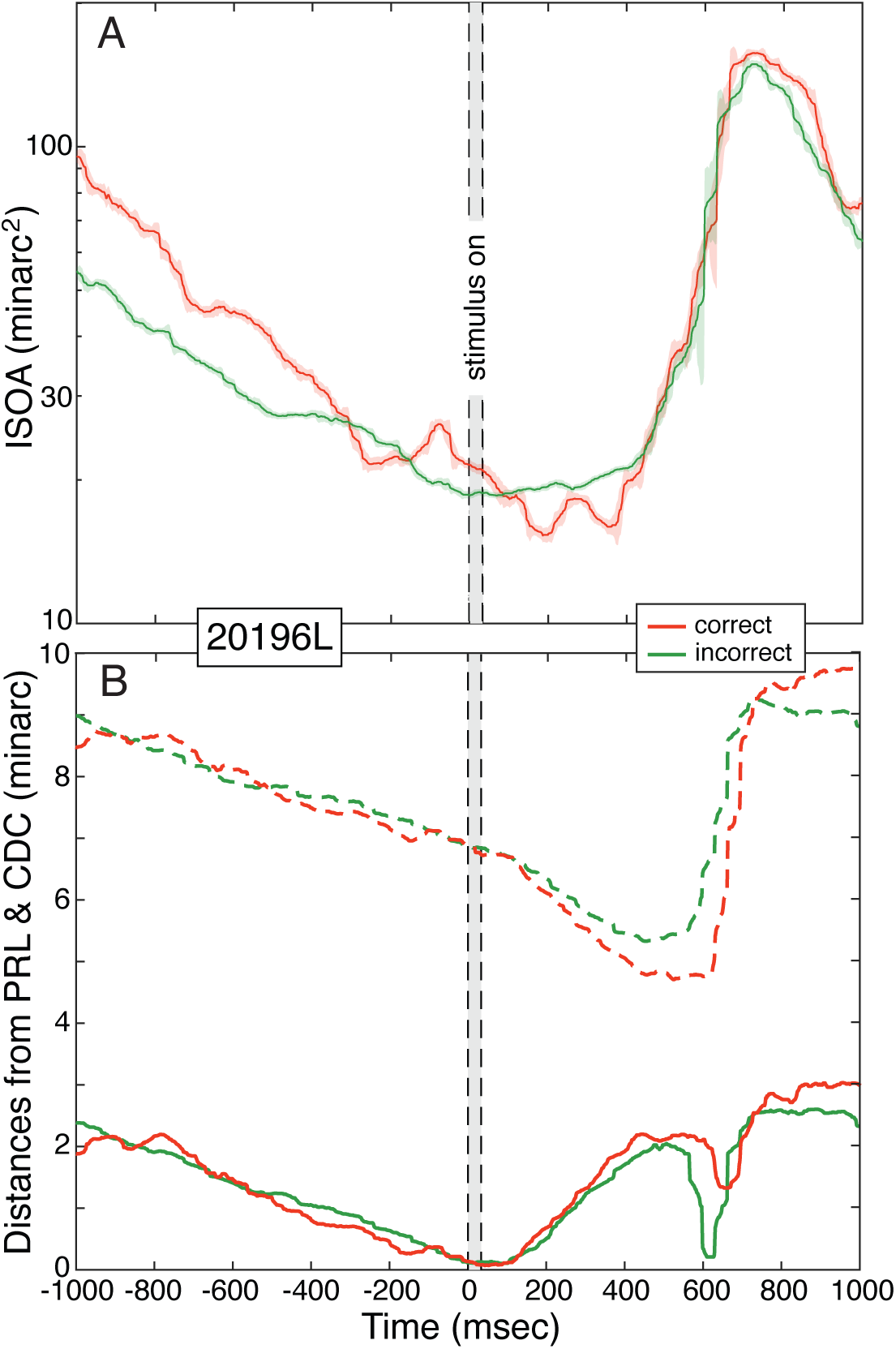
Variation of fixation during the course of a trial. **A)** ISOA over time. Area of the ISO contour containing 68% of retinal positions is plotted over the 2-sec duration of a trial for a representative subject. The time at which the stimulus was presented is represented by the gray bar. The data have been averaged across all 2100 trials. A running median was applied at each time point over 20 preceding and 20 succeeding time samples. Green and red lines represent contour areas on correct and incorrect trials, respectively. Shaded regions around each line indicate standard deviations. **B)** Fixation relative to PRL and CDC during the course of a trial. The lower solid curves represent median distance of fixation from the PRL over the duration of a trial for the same subject. The distance goes to zero at the time of the stimulus, by definition. The gray bar represents stimulus presentation time. The data have been averaged across 2100 trials. The green and red lines represent distance from PRL on correct and incorrect trials, respectively. The upper dashed curves represent the median distance of fixation from the CDC for the same subject. Again green and red represent correct and incorrect responses.

Another indication of the consistency of fixation is illustrated by Fig. 4B, which plots distances from the PRL and CDC over the course of trials for the same subject. The solid lines represent distance from the PRL. As stimulus presentation approached, fixations became systematically closer to the PRL and remained close as the stimulus was presented and then for a few hundred milliseconds after the stimulus was extinguished. The dashed lines represent distance from the CDC. Again as the stimulus time approached, fixation became closer to the CDC, but never as close as they were to the PRL. As shown in Fig. 2A, the PRL was between the saccade landing point and CDC, so movement toward the PRL also decreased distance to the CDC. The other subjects exhibited similar behavior (see Fig. S4).

We computed the median trajectories of the eye over the course of the 2-sec trials. Fig. 5 plots the trajectory for a representative subject. The 3D plot has time flowing from the top to the bottom. The green and red traces are the trajectories on trials where the subject’s response was correct and incorrect, respectively. The orange column shows the position of the fPRL. The plot illustrates the consistency of fixational eye movements when performing a visual-resolution task. At the beginning of a trial, this subject fixates above and to the left of where the stimulus will appear. As stimulus presentation approaches, consistent drift down and to the right occurs; the drift brings the fPRL toward where the stimulus will appear. The drift continues in the same direction as the stimulus is presented and for a couple hundred milliseconds after the stimulus has been extinguished. The substantial decrease in trajectory variability (represented by the transparent green and red disks) during the drift shows that the eye’s trajectory during the drift is much more consistent than its trajectory near the start and end of the trial. The variability of the drift appears to be slightly less for correct than for incorrect ones. This observation is supported by the fact that ISOA is smaller at the time of stimulus presentation for correct than incorrect trials (Figs. 4 & S3). Fig. S5 shows that every subject exhibited this general pattern: consistent downward drift before, during, and after stimulus presentation. The horizontal component of the drift, however, varies from subject to subject. The trajectories near the end of a trial and the beginning of the next one were dominated by microsaccades and were less consistent than the drifts.

**Figure 5:**
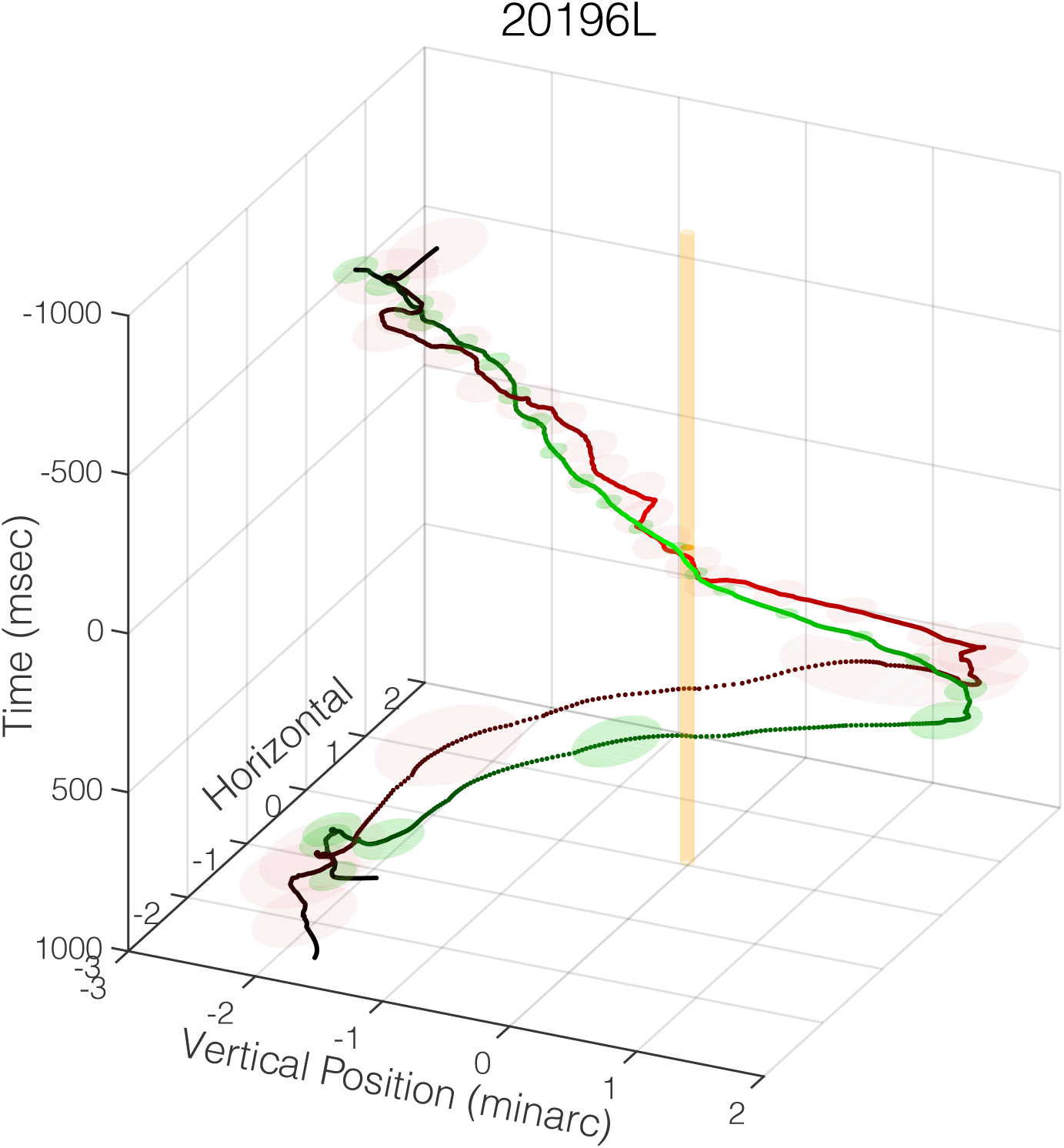
Median eye trajectories over the course of the 2-sec trials for a representative subject. The 3D coordinates are horizontal position and vertical position in minarc and time in msec. A 3D coordinate was calculated for every time sample in each trial. The median of those points was computed for each time sample and contours were drawn from those points using a Savitzky-Golay filter to smooth the data. The green and red traces are the trajectories on correct and incorrect trials, respectively. The trajectories start at the top, 1000msec before stimulus appearance, and end at the bottom, 1000msec after. The transparent green and red ellipses represent standard errors every 76msec. Trials in which the Vernier gap was +/-18secarc have been excluded because performance was at ceiling on those trials. Including them does not, however, affect these trajectories noticeably. The orange cylinder represents the fPRL. An animation of the 3D plot is shown in Supplemental Materials Movie S1.

## Discussion

### Fixation Strategy & Acuity

Humans adopt a machine-like fixation strategy that places an anticipated stimulus on a very specific location of the retina. This locus, which we call the functional PRL, is significantly smaller than the overall PRL (Table 1, Fig. 2A). Recent studies using retinal imaging and stabilized stimuli have reported that visual sensitivity and visual acuity are no better at the PRL than at the anatomical fovea or CDC (*30, 31*). We re-examined this by using a unique methodology to look for changes in performance near the center of the fovea. First, we chose a hyperacuity task—Vernier acuity—which is more affected by retinal eccentricity than other visual tasks such as letter acuity (*22*). Second, we allowed subjects to make natural fixational eye movements during a cadenced task, which enabled them to position the stimulus and its motion on the retina in a way that presumably optimized performance.

We observed a measureable drop in acuity as the stimulus landed farther from the fPRL, but the drop was small and did not reach statistical significance until the stimulus was more than 7–10minarc from that location (Fig. 2C). Thus, acuity was not significantly better at the fPRL than at the CDC (*i.e.*, center of the anatomical fovea).

Fixation may be considered a voluntary task, but the properties of fixation in a given task are not (*44*). We are generally unaware of our drifts and microsaccades, so we do not guide fixation via conscious awareness of when fixation deviates. Yet our data show that fixation is sufficiently good to maintain a high and consistent level of performance in a demanding task. On only 17% of the trials did the fixational movements lead to a small but consistent drop in performance in the Vernier-acuity task. This suggests that control of fixation is more precise than a subject’s ability to perceive an error in fixation. This point is supported by an earlier study that found that humans’ awareness of the direction of gaze was more than 40% less precise than the actual spread of fixation (*26*). It is sensible for the oculomotor system to enable a narrower spread of fixation than conscious behavior reveals. Consider an alternative situation in which fixation was guided by conscious awareness of gaze direction. If this occurred, performance in the acuity task would have been worse than we observed.

Our observation of more precise oculomotor behavior than perceptual behavior is analogous to the findings that people make accommodative responses to changes in retinal-image blur that are not perceived (*45*) and make vergence eye movements to binocular disparities that are not perceived (*46*). Thus, oculomotor behavior that is more precise than perceptual awareness may be a general principle.

### Role of Microsaccades

Microsaccades are thought to redirect gaze toward the intended fixation target (*47*). We find on a fine scale that they do not. Two observations support this claim. 1) The ISOA for microsaccade landing points is about three times larger than the fPRL (Fig. 2A). This means that the drift that follows a microsaccade also serves to direct gaze toward the fPRL. This point is illustrated by the trajectories in Figs. 5 and S5 that show drifts in a fairly constant (but idiosyncratic) direction as the stimulus approaches, is presented, and then extinguished. 2) Most subjects exhibit a form of sub-clinical micronystagmus during fixation (*28*). Inter-saccadic drifts generally have a downward tendency in gaze direction. The oculomotor system takes this into account by making upward microsaccades that direct the anticipated stimulus slightly above the fPRL rather than toward it. This behavior is evident in Figs. 5 and S5.

If the end point of the microsaccades placed the stimulus in a location that was more optimal for vision, we would have observed lower Vernier-acuity thresholds in the vicinity of the saccade landing points; we did not.

### PRL vs. Anatomical Fovea

The fPRL was generally slightly above the location of the anatomical fovea, or CDC, in fundus coordinates, consistent with previous reports (*24, 25, 28*).

It has been reported that drift movements tend to redirect images toward the CDC (*32*). If the region of greatest cone density actually conferred a functional advantage, it stands to reason that the functional PRL would be centered at that location: but it is not. In fact, in all subjects the stimulus rarely landed at the CDC. Figs. 4B and S4 show that drifts do indeed tend to be toward the CDC, but they never bring the stimulus location closer to the CDC than to the fPRL before, during, or after stimulus presentation. The observation that drifts tend toward the CDC is instead a consequence of the fact that the fPRL lies between saccadic landing point and the CDC (Fig. 2A). Thus a movement toward the fPRL is also a movement toward the CDC.

Reiniger and colleagues (*25*) suggested that the upward displacement of the PRL (downward in world coordinates) relative to the locus of CDC is advantageous for vision in the natural environment. They pointed out that scene elements above the current fixation point are generally farther than the fixated element, and that the elements below fixation are generally nearer (*48, 49*). The vertical displacement of the PRL from the CDC means that scene elements above the direction of gaze are imaged onto the CDC. Reiniger and colleagues claimed that more distant elements create more power at high spatial frequencies than near elements and that having the region of greatest cone density directed toward those more distant parts enables more accurate measurement of image features there. Unfortunately, there are two problems with this hypothesis.

First, farther parts of natural scenes do not have greater power in high spatial frequencies than nearer parts. The power spectra of natural images tend to fall with the square of spatial frequency: 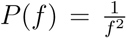, where *f* is spatial frequency and *P* (*f*) is power as a function of frequency (*50–52*). Images with such power spectra are scale-invariant; *i.e.*, their statistical properties do not change if one changes the scale at which the images are observed (*50*). Thus, this part of the hypothesis of Reiniger and colleagues is not true for images created by the natural environment.

Second, we observed that acuity—a measure of sensitivity at high spatial frequencies—is actually no better at the CDC than at the PRL. So the greater receptor density at the peak does not confer a resolution advantage compared to the retina at the PRL. (Note that the increased cone density at the CDC compared to the PRL is only expected to confer at 2% improvement at best; Table 1.) Therefore, this part of the hypothesis is not supported by our data.

Thus questions remain.

First, why is the PRL generally displaced from the CDC? When we began this project we thought that we might find that acuity is better at the PRL than in the region of highest cone density. If this were true, it would help us understand why fixations are directed toward the small and consistent PRL. But we did not observe an increase at the PRL compared to the CDC, so our data do not provide an answer to the question of why the PRL is generally displaced from the region of highest cone density.

Second, why is the PRL usually displaced upward from the center of the fovea in retinal coordinates? We do not have a persuasive answer, but it is interesting to note that the PRL in patients with central scotomas is usually also displaced upward in retinal coordinates from the center of the fovea (*53–55*). Perhaps whatever drives the visual system to have an upward displacement in people with compromised vision is the same as what drives a small upward displacement in those with normal vision.

### Are Human Fixational Eye Movements Optimal?

There is no simple answer to this question.

In support of the idea that fixational movements aid performance, we find that microsaccades and drift work in concert to place an anticipated stimulus within a small retinal region, and that performance at or near this fPRL is better than in other nearby regions (Fig. 2C). The spatial extent of fixational eye movements is so small that the cost of fixation instability has little consequence for visual performance. Specifically, in our demanding visual task, fixation instability led to only a 34% drop in performance on only 17% of the trials (Fig. 2C). This exquisite control of gaze is executed without conscious awareness.

We also find that the timing of microsaccades matters. When they occur too close in time to the onset of the stimulus, performance suffers (Fig. 3B).

But some of our observations seem inconsistent with the idea that fixational eye movements optimize performance. For example, the greatest suppression of microsaccades occurs after the stimulus has come and gone (Figs. 3A & S2), and this seems non-optimal. We note, however, that during natural viewing, humans make roughly two saccades or microsaccades per second (*56,57*), which means that stimuli remain in a fairly constant retinal position for a bit less than 500msec. Perhaps our observation of maximum suppression 200-400msec after stimulus presentation reflects an oculomotor strategy to suppress eye movements for the duration of a typical natural fixation. It is interesting to note as well that the time required to maximize visual acuity is about 400msec (*58*).

The fact that the fPRL is displaced from the region of peak cone density (CDC) also seems non-optimal. But the drop in linear cone density from the CDC to the fPRL is only 2% (Table 1), so the expected loss in effective resolution due to fixating slightly off the CDC is trivial. Moreover, the mapping between the central retina and higher visual areas, such as visual cortex, is strongly affected by the post-natal migration of cones toward the retinal center and the migration of retinal neurons away from that region (*59,60*). It is thus perhaps not surprising that the retinal region for preferred fixation in adults is not identical to the region of highest cone density. Indeed, it is remarkable that they are so close to one another.

## Materials and Methods

### Subject recruitment

Six subjects participated. All self-reported to have normal vision and no ocular disease. Data from one subject were dropped after learning that they initially misunderstood how to respond in the Vernier task. Of the remaining subjects, average age was 28.2 years. Three were female. Two (20196 and 20109) were authors. Cycloplegia and mydriasis were induced with drops of 1.0% tropicamide and 2.5% phenylephrine hydrochloride ophthalmic solution. All procedures were approved by the Institutional Review Board at the University of California, Berkeley. Informed consent was obtained from each subject before the experiments.

### AOSLO system

The AOSLO is a custom-built, scanning-laser device that can image the retina while also delivering images to the retina. The system uses adaptive optics to measure and correct for optical imperfections that would otherwise blur the images. More details on the system are published elsewhere (*61, 62*). Only details that are directly relevant to this study are described here.

The AOSLO uses different wavelengths for imaging and psychophysics. We used 940-nm light for wavefront sensing and 680-nm light for imaging and stimulus delivery. The scanned field size was 0.9×0.9^◦^ and each frame was digitized into 512×512 image pixels yielding a pixel size of 6.33secarc. Video imaging was done at about 30 frames per second.

The raster scan appeared to the subject as a very bright bright red square field (estimated to be 5.8 log Trolands (*63*)). A set of fixation guides at the corners of a 25×12.5minarc rectangle were present in every frame (Fig. 1C). For two frames out of every 60-frame sequence (2sec), a Vernier stimulus consisting of two horizontal bars appeared at the center of the fixation guides. The bars were 0.95×1.9minarc (9×18 pixels) and the horizontal gap between them was 0.95minarc. The fixation guides and Vernier stimulus were presented within this field by turning off the scanning laser at specific time points during the scan (*20, 64*). The same laser that was used for imaging was used to present the decrement stimulus, so the actual stimuli appeared as dark pixels in the recorded image of the retina. This offers an unambiguous record of the exact location where the stimulus landed, with cellular-level accuracy. The stimulus appeared ultra-sharp to the subject owing to the use of adaptive optics and large pupil (7.2mm).

### Vernier-acuity measurement

We used the method of constant stimuli to estimate the smallest gap yielding reliably correct performance. Upward and downward offsets of 0, 1, 2 and 3 pixels (0, 6.33, 12.66, and 18.99secarc) between the two bars were presented in pseudo-random order. Data were collected for blocks of 21 trials over the course of a 44-sec AOSLO video. 100 blocks were conducted for each subject. The stimulus was delivered for two frames at the middle of each 2-sec trial. In a 2-alternative, forced-choice judgment, the subject indicated on each trial whether the left bar had been higher or lower than the right bar. No feedback as to the correctness of the response was provided.

Threshold was estimated by fitting the resulting psychometric data with a logistic function using a maximum-likelihood criterion (*42*). From the fitted function, we found the positive (left bar up) gap size yielding 75% left responses and the negative (left bar down) gap size yielding 25% left responses. The threshold estimate was the difference in those two sizes divided by two. We also computed 95% confidence intervals from the psychometric fits.

### Image- & eye-motion analysis

Incessant eye movements that occur during AOSLO scanning caused unique distortions within each frame of the recorded video. We used versions of previously published algorithms to remove distortions and stabilize each recorded video from each subject (*65*). Correcting for these distortions served two purposes.

First, it enabled the summing of multiple frames to provide high signal-to-noise images. All frames in each distortion-corrected video were added to generate a high-quality image of the foveal cone mosaic. The smallest cones in the center of the fovea could be resolved in all of our subjects. The best-appearing image from one of the videos for each subject was used as their master retinal image. In this image semi-automated methods were used to label every cone (*66*). Cone-density maps were computed by integrating all the labelled cones within a 10minarc circular area around each pixel in the master image (*24*). The cone density centroid (CDC) was computed as the density-weighted centroid within the contour at 80% of the peak density, as per methods described by (*25*). An example master retinal image with cone density colormap overlay from one subject is shown in Fig. 1B.

Second, the byproduct of the distortion correction is a high-speed, high-accuracy eye-motion trace. For the videos, we corrected the distortion at 32 segmented strips across each scanned frame and generated eye-motion estimates at 960Hz. Each eye-motion trace from each video was annotated for drifts, microsaccades, and blinks using semi-automated, custom software. Segments with bad tracking due to poor image quality were dropped from each trace, but there were few instances and of those they were short in duration.

The eye-movement traces indicate how a continuously viewed scene would have moved across the retina during fixation. To make each eye-motion trace yield the exact retinal trajectory of the Vernier stimulus, we added a fixed offset to the trace to align its *XY* locations with the *XY* location of the stimulus on the frames in which it appeared. Fig. 1D shows a master retinal image with 960-Hz traces overlaid for a single video. The orange segments of the trace indicate when the stimulus appeared. Contour plots of the entire trace or selected elements of it (*e.g.*, saccade start points, saccade end points) and corresponding ISOA contours were generated and plotted on the same master retinal image from all trials. Despite training, some subjects landed their saccade more than three standard deviations away from the others, probably in an attempt to search for the stimulus when they did not perceive it. Such landing positions were rejected as outliers and not included in the computation of the ISOA contours.

## Acknowledgments

We thank Pavan Tiruveedhula for technical support on this project.

## Author Contributions

Conceptualization: AR, MB. Methodology: AR, MB, JG, NB. Investigation: AR, MB, JG, NB. Formal analysis: AR, MB, JG, NB. Resources: AR. Funding acquisition: AR. Project administration: AR, MB. Supervision: AR, MB. Visualization: AR, MB, JG, NB. Data curation: AR, JG, NB. Validation: AR, MB, JG, NB. Software: AR, MB, JG, NB. Writing—original draft: AR, MB, JG, NB. Writing—review and editing: AR, MB, JG, NB.

## Data and Materials Availability

All data needed to evaluate the conclusions in the paper are present in the paper and/or the Supplementary Materials.

## Supplementary Materials for

**Movie S1.** Animation of 3D plot from Fig. 5

**Movie S2.** Animation of 3D plot from Fig. S5

**Figure S1:**
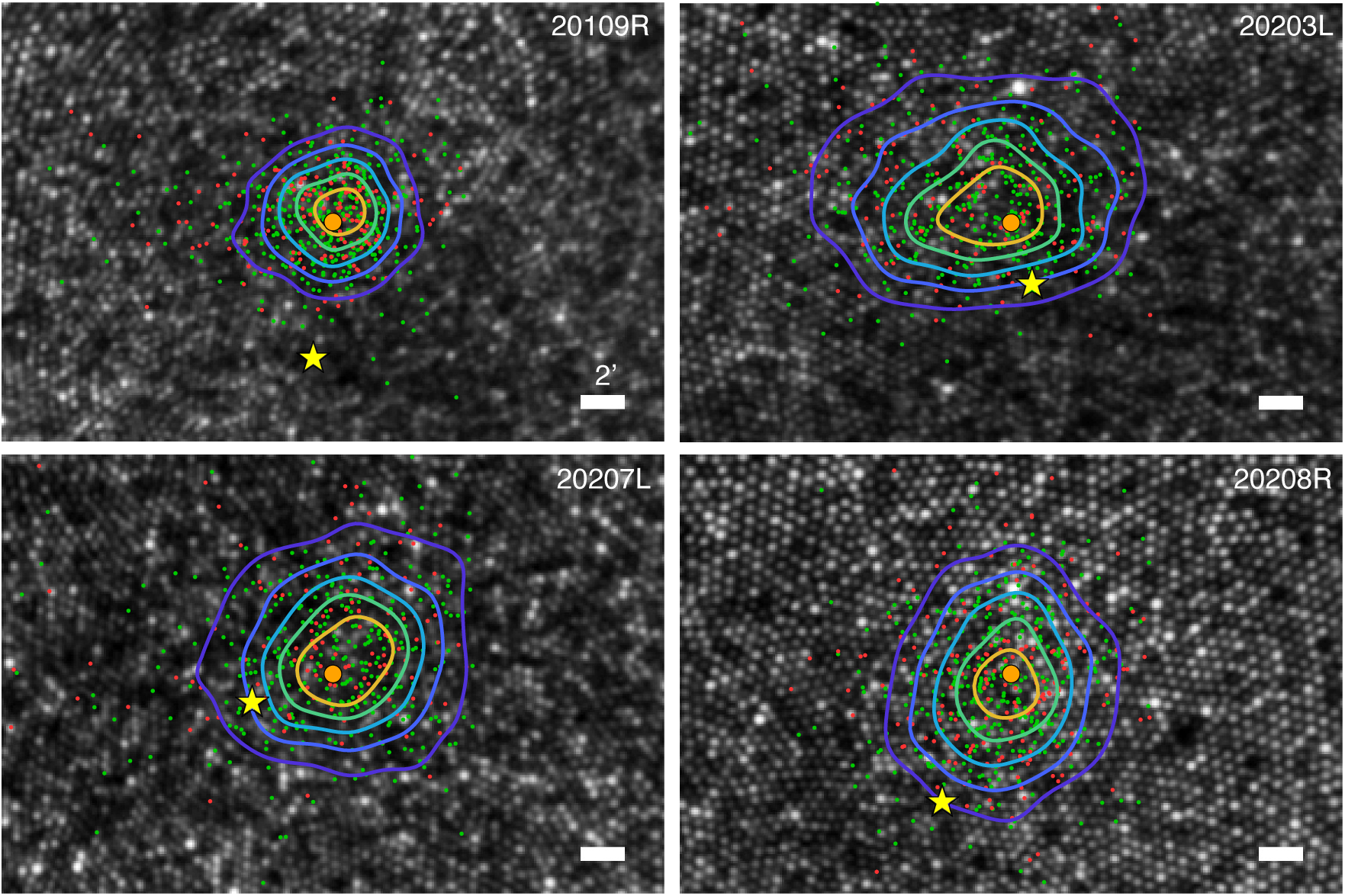
Stimulus positions and performance for the four subjects not shown in Fig. 2B. The position of the stimulus is plotted for each subject. Green and red dots represent those positions on trials in which the response was correct and incorrect, respectively. Orange circles represent the PRL. Yellow stars represent the cone-density centroids (CDCs). The colored contours indicate, respectively, 17% of positions falling closed to the PRL (yellow), 18–33% (green), 34–50% (cyan), 51–67% (blue), and 68–83% (purple).

**Figure S2:**
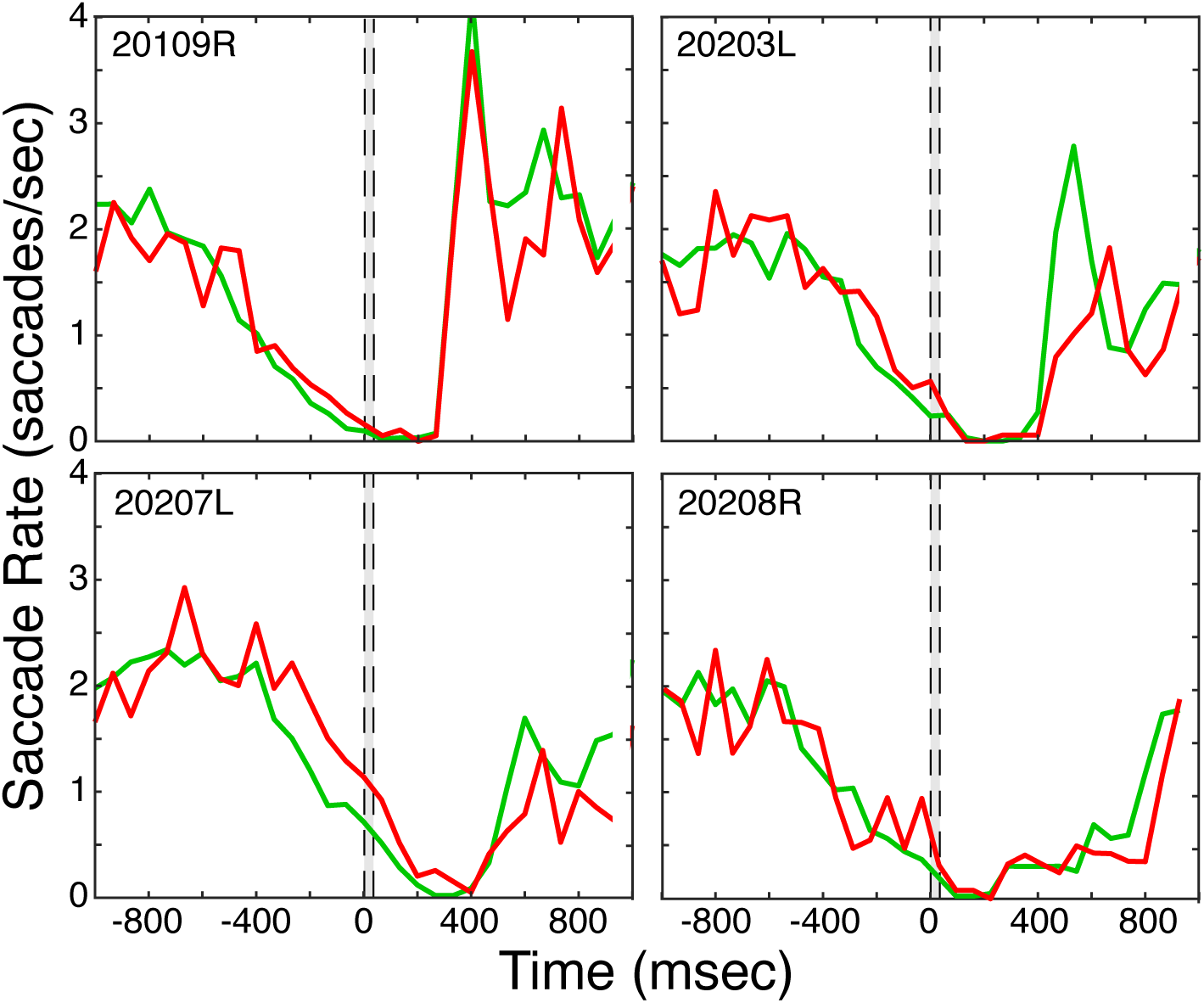
Saccade rate over time. The average number of saccades per second is plotted as a function of time for the four subjects not shown in Fig. 3A. Stimulus onset is at time 0. The green and red lines represent saccade rate during trials in which the observer responded correctly and incorrectly.

**Figure S3:**
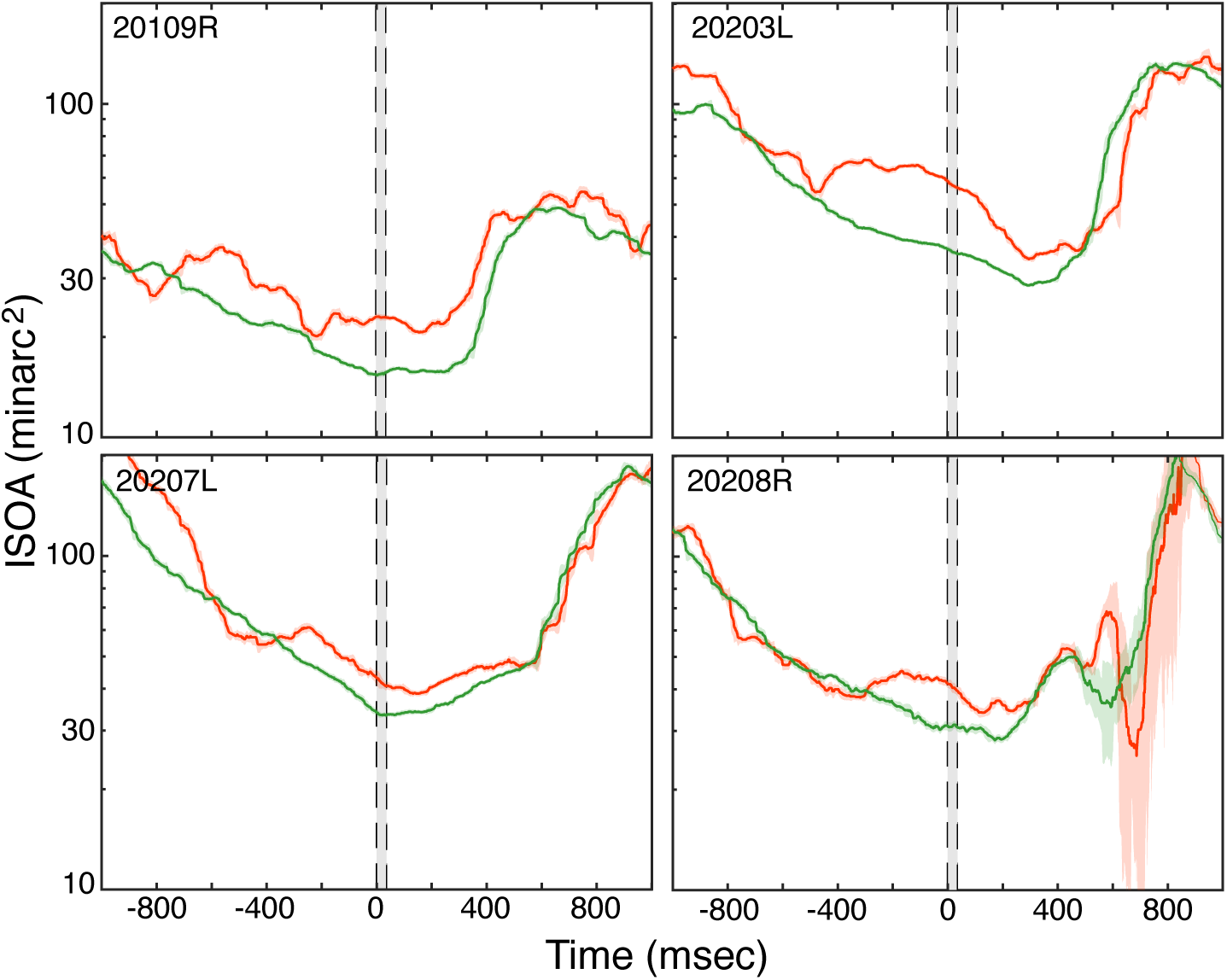
ISO area over time for the four subjects not shown in Fig. 4A. The area of the ISO contour containing 68% of retinal positions is plotted over the 2sec duration of a trial. The time at which the stimulus was presented is represented by the gray bar. The data have been averaged across the 2100 trials the subjects encountered. For each time a running median was computer over the 20 preceding and 20 succeeding time samples. The green and red lines represent contour areas on correct and incorrect trials, respectively. The shaded regions around each line indicate standard deviations.

**Figure S4:**
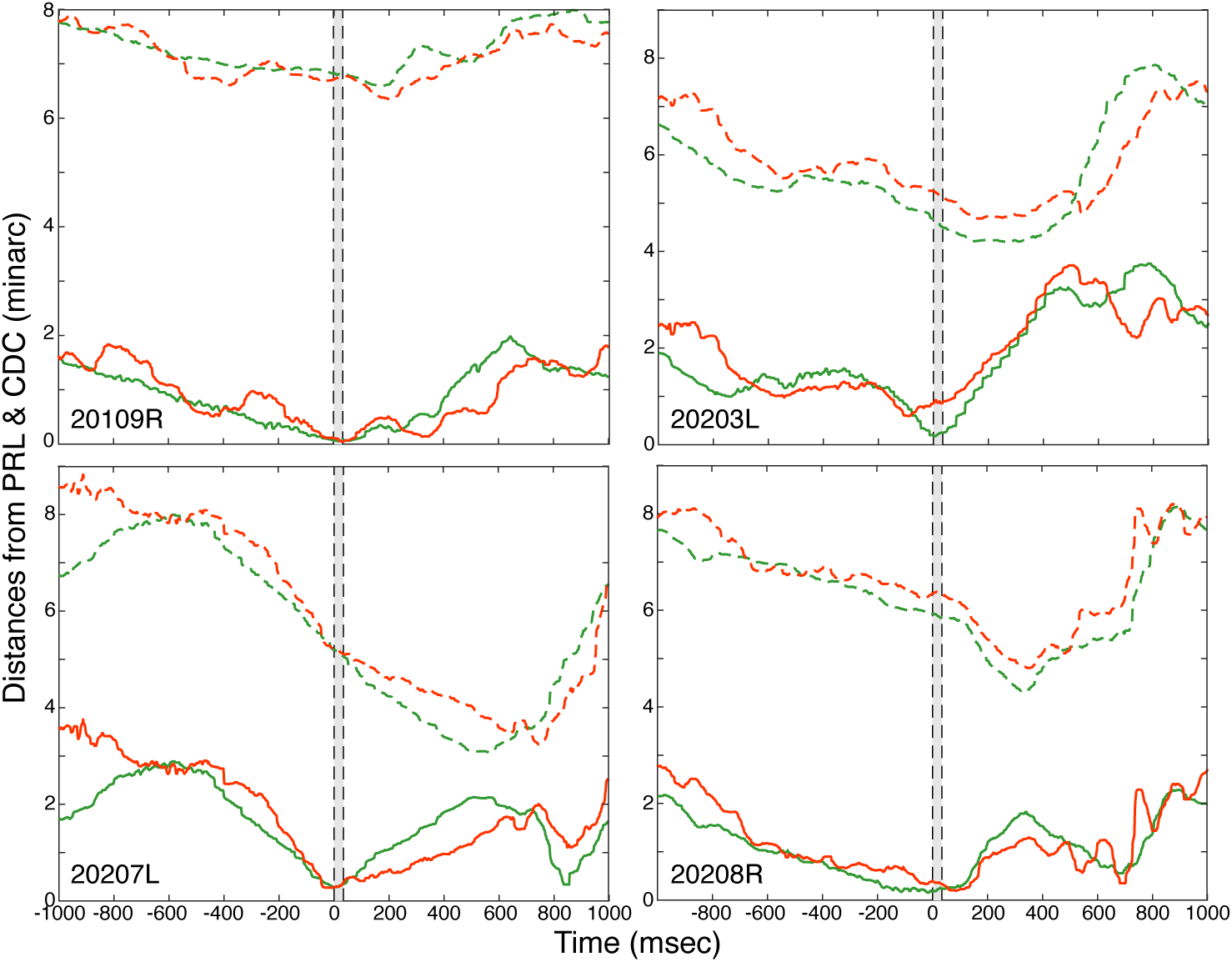
Fixation relative to PRL and CDC during the course of a trial for the four subjects not shown in Fig. 4B. The solid lower contours represent the median distance of fixation from the PRL the duration of a trial. The time at which the stimulus was presented is represented by the gray bar. The data have been averaged across the 2100 trials the observers encountered. The green and red lines represent distance from PRL on correct and incorrect trials, respectively. The dashed upper contours represent the median distance of fixation from the CDC. Again green and red represent distance on correct and incorrect trials, respectively.

**Figure S5:**
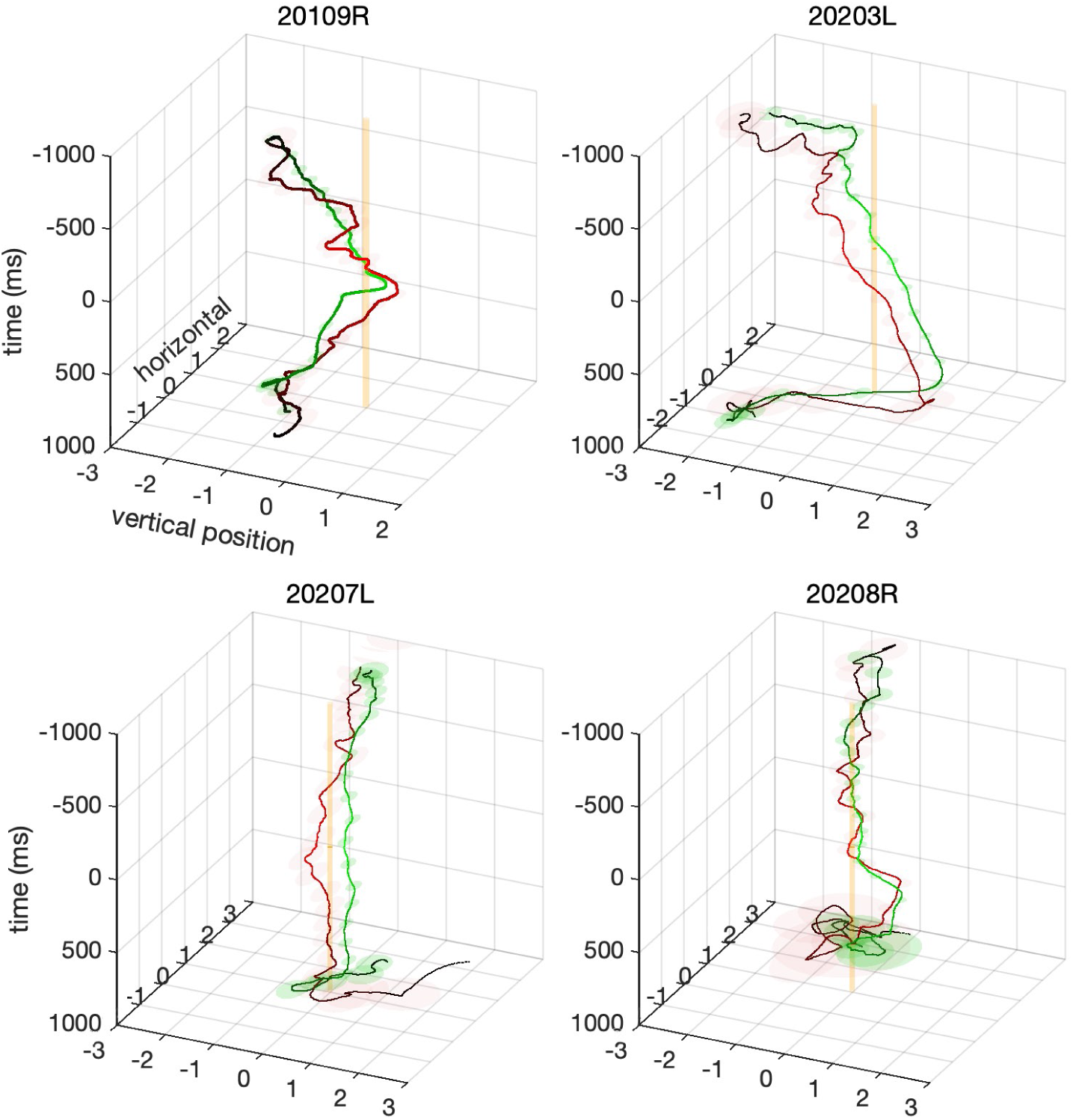
Median eye trajectories over the course of the 2-sec trials for the four subjects not shown in Fig. 5. The 3D coordinates are horizontal position and vertical position in minarc and time in msec. A 3D coordinate was calculated for every time sample in each trial. The median of those points was computed for each time sample and contours were drawn from those points using a Savitzky-Golay filer to smooth the data. The green and red traces are the trajectories on correct and incorrect trials, respectively. The transparent green and red ellipses represent horizontal and vertical position variability every 76msec. Trials in which the Vernier gap was +18secarc have been excluded because performance was at ceiling on those trials. Including them does not, however, affect these trajectories noticeably. The orange cylinders represent the fPRL. An animation of the 3D plot is shown in Supplemental Materials Movie S2.

